# Feature distribution learning by passive exposure

**DOI:** 10.1101/2021.10.13.464193

**Authors:** David Pascucci, Gizay Ceylan, Árni Kristjánsson

## Abstract

Humans can rapidly estimate the statistical properties of groups of stimuli, including their average and variability. But recent studies of so-called *Feature Distribution Learning* (FDL) have shown that observers can quickly learn even more complex aspects of feature distributions. In FDL, observers learn the full shape of a distribution of features in a set of distractor stimuli and use this information to improve visual search: response times (RT) are slowed if the target feature lies inside the previous distractor distribution, and the RT patterns closely reflect the distribution shape. FDL requires only a few trials and is markedly sensitive to different distribution types. It is unknown, however, whether our perceptual system encodes feature distributions automatically and by passive exposure, or whether this learning requires active engagement with the stimuli. In two experiments, we sought to answer this question. During an initial exposure stage, participants passively viewed a display of 36 lines that included one orientation singleton or no singletons. In the following search display, they had to find an oddly oriented target. The orientations of the lines were determined either by a Gaussian or a uniform distribution. We found evidence for FDL only when the passive trials contained an orientation singleton. Under these conditions, RT’s decreased as a function of the orientation distance between the target and the exposed distractor distribution. These results suggest that FDL can occur by passive exposure, but only if an orientation singleton appears during exposure to the distribution.

## Introduction

Humans can extract meaningful information from complex visual scenes in a fraction of a second. Most of the information that is available in the immediate ‘gist’, takes advantage of statistical regularities and redundancies typically found in the visual world. Similar objects are often arranged into groups defined by distributions of low-level features, like color, orientation, size, and location. This provides the visual system with the opportunity to form coarse, global representations, without recognizing individual details: “there is a big basket of small red apples on the left side of the grocery shop”. But how coarse are these global representations that are automatically and effortlessly available?

A large body of evidence indicates that humans can rapidly extract basic statistical summaries, like the average and variability of an ensemble of similar stimuli (Ariely, 2001; Whitney & Leib, 2018). Recent studies, however, demonstrate that the perceptual system can quickly learn even more complex and detailed aspects of stimulus distributions, such as the whole shape of a distribution of visual features. In studies of *feature-distribution learning* (FDL; Chetverikov et al., 2016, 2017b, 2017a), for instance, observers are presented with a sequence of visual search trials containing a singleton target embedded in a set of distractors (see Figure 1). After only a few exemplars of distractors sharing a similar distribution of features, visual attention learns the shape of the distribution and uses this information to guide future search: searching for a new target becomes easier if the target lies outside the distribution of previous distractors, and the response time functions reflect the shape and characteristics of the distractor distribution. This suggests that even when ensembles are not directly relevant for behavior (e.g., a set of distractors in a search task), the brain encodes detailed representations of their properties, beyond summary statistics, which can ultimately aid the detection of task-relevant outliers. This form of learning has been observed for both colors (Chetverikov et al., 2017b) and orientation (Chetverikov et al., 2016), for various distributions types (even bimodal ones, Chetverikov et al., 2017a) and occurs implicitly, since observers cannot explicitly judge distribution shapes, even though the learning strongly affects their search performance (Hansmann-Roth et al., 2021).

**Figure 1.**
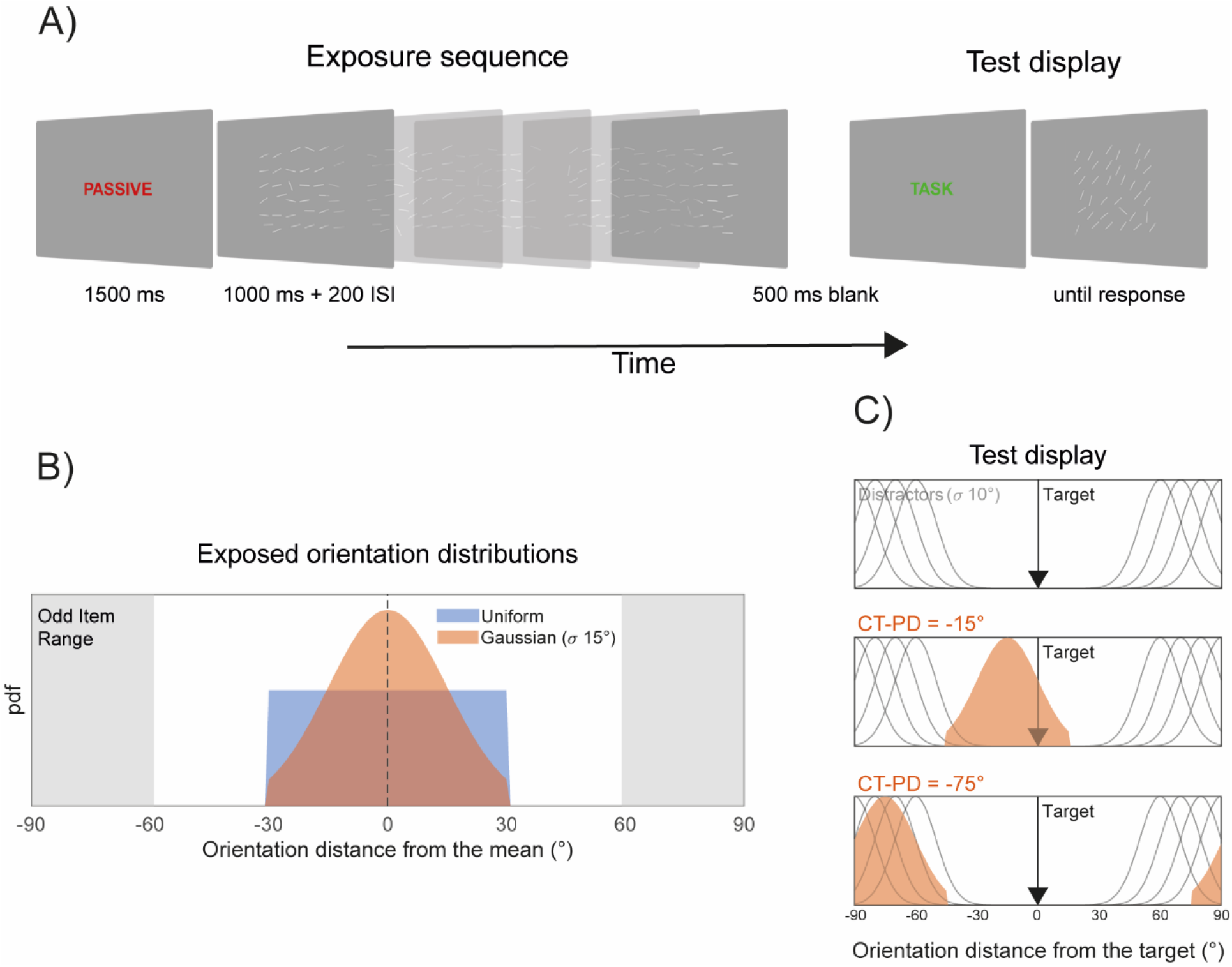
Trial structure and main variables. A) Example of a sequence of displays in one trial. Participants were instructed by a written cue to passively view the display (exposure sequence) or to perform a visual search (test display). In the visual search task, they had to report the location of the oddly oriented target (upper vs. lower quadrant). B) In the exposure sequence, the orientations in the ensemble of lines were drawn according to two different distributions, a truncated Gaussian, and a uniform distribution. One oddly oriented line with orientation distance of ± 60-90° from the ensemble mean, was included in the exposed displays of Experiment 1, but not in Experiment 2. C) In the test display, distractors followed a Gaussian distribution with 10° of standard deviation and a mean located between ±60° and ±90° away from the target orientation (gray curves). The target orientation was presented at different distances from the mean of the exposed distractors distribution (current-target previous-distractor distances, CT-PD; orange Gaussian distribution in the example).

An important question concerning this learning is whether such detailed statistical representations are encoded automatically and independently of a task —i.e., by simple exposure. In FDL, observers are engaged in a search task that requires them to segment out a potential target from a set of task-irrelevant features. In most of our everyday routines, however, we are not constantly engaged in visual search. Yet, learning the distribution of visual features can help to maintain an accurate representation of our visual environment while maximizing our ability to identify outliers and relevant changes in the environment. In line with this, research has shown that our attentional and perceptual systems constantly learn the characteristics of the visual input, even while idling, giving rise to phenomena of latent learning and plasticity (Turatto et al., 2018; Won & Geng, 2020). In studies of the habituation of the attentional capture response, for example, simple exposure to a repetitive onset stimulus can prepare the attentional system for resisting capture by the same onset in the context of a task (Turatto et al., 2017, 2018; Turatto & Pascucci, 2016).

While feature distribution learning from active search has been found to aid visual performance, this mechanism would be more useful if it can also operate more generally, outside this task-specific context. In the present work, we, therefore, asked whether passive exposure to sets of oriented lines coming from distributions of orientations can induce feature distribution learning effects that modulate visual search during a subsequent active search task. We presented a sequence of five displays containing 36 oriented lines. All the lines were oriented according to either a Gaussian or a uniform distribution (see Methods and Figure 1), except an oddly oriented one resembling the target of a typical FDL display. In a sequence of short blocks, observers were asked to passively view each display without performing any explicit task. After the sequence, a single *test* display was presented, requiring observers to perform a search task, reporting the location of the singleton target (upper vs. lower quadrant). In the test display, we used *role-reversal* effects in visual search (Kristjánsson & Driver, 2008) to probe signatures of FDL: search times were analyzed as a function of the angular distance between the mean of the prior distractor distribution and the orientation of the test target (the Current-Target/Previous-Distractor distance, CT-PD) (Chetverikov et al., 2019).

## Methods

### Participants

A total of 40 healthy participants (24 in Experiment 1 and 16 in Experiment 2, age range of 19-35 years, 18 females), from the EPFL and the University of Lausanne, participated in the study for a monetary reward (20 CHF/hour). All participants had normal or corrected-to-normal vision and were naïve as to the purpose of the experiments. Before the experiments, visual acuity was tested with the Freiburg Acuity test (Bach, 1996). A value of 1 or above reached with both eyes open was used as the selection criterion. The sample sizes were selected based on previous FDL studies. All participants gave written informed consent, and the study was approved by the local ethics committee under the Declaration of Helsinki (apart from preregistration) (World Medical Organization, 2013).

### Apparatus

Stimuli were presented on a gamma-corrected VG248QE monitor (resolution: 1920 x 1080 pixels, refresh rate: 120 Hz) and were generated with custom-made scripts written in Matlab (R2013a) and the Psychophysics Toolbox (Brainard & Vision, 1997), running on a Windows-based machine. Experiments were performed in a darkened room and participants sat at 57 cm from the computer screen, with their head positioned on a chin rest.

### Stimuli and procedure

In both experiments, trials were arranged into short mini-blocks comprising an exposure sequence of five displays (1000 ms each, separated by a 200 ms blank screen), followed by a single test display (see Figure 1). All stimuli were presented on a grey background. A colored written cue (1500 ms) indicated the upcoming sequence of exposed displays (the word ‘PASSIVE’, written in red) or the test display (‘TASK’ written in green). Each display contained 36 white lines of 1° length, arranged in a 6-by-6 array within a square of 14° centered at the fovea. A small jitter was added to the location of each line within each cell of the 6-by-6 array (jitter range: ±0.5°) to prevent uniform appearance.

During the exposure stage, the orientation of each line was drawn from a truncated Gaussian distribution (standard deviation (σ): 15°, cut-off at ± 30° [2σ] from the mean) or a uniform distribution (range: ± 30° from the mean), in separate and intermixed mini-blocks consisting of 5 learning trials and a test trial. The mean orientation of both distributions was randomly selected for each mini-block. In Experiment 1, the exposure displays always contained a singleton line, oriented between ±60° and ±90° away from the mean of the distribution. In Experiment 2, no singleton line was present in the passively viewed displays during the exposure sequence.

On test trials, the orientation of the target line was drawn from among a set of 12 angular distances (from −75° to +75° in steps of 15°) from the mean of the previous sequence of exposed distributions. The distance between the target line orientation and the previous distributions mean defined the Current-Target/Previous-Distractor distance (CT-PD). The distractor distribution of orientations in the test display was always Gaussian with σ of 10° and the mean located between ±60° and ±90° away from the target orientation.

The target line in the test display could be presented in one of the 36 possible locations. Participants were asked to report the location (upper vs. lower quadrant) of the target line by pressing the ‘i’ (upper) or ‘j’ (lower) keys of a computer keyboard as quickly as possible. The test display remained on the screen until a response was made. Two types of feedback were used to ensure that participants engaged in the task and maintained a high accuracy rate. First, error feedback (the word ‘error’, written in black) was presented following incorrect responses. Second, a score was calculated on each mini-block (as in previous work, Chetverikov et al., 2019), and the average score was shown after every 33 mini-blocks.

At the beginning of each experiment, participants were provided with written and verbal instructions and performed a sequence of practice trials under the supervision of the experimenter.

Practice trials were not analyzed further but served to ensure that participants understood the task. Experiments consisted of 264 mini-blocks with 8 breaks, lasting approximately 1 hour.

### Analysis

Trials with errors and response times larger than 1 second were excluded from the analysis, following guidelines from previous work (Chetverikov et al., 2019) (34.6% in total for Experiment 1, 23.8% for Experiment 2). Participants performed with an average accuracy of 94±4% in Experiment 1 and 93±4% in Experiment 2. In Experiment 2, one participant was excluded due to average response times larger than 2 seconds, well beyond the threshold recommended in previous FDL studies (Chetverikov et al., 2019).

To evaluate the effect of the exposed distributions on search times, the relationship between single-trial log-transformed response times and the CT-PD variable was modeled with a set of 5 models, which included a model assuming no effect of CT-PD (‘constant’) and models assuming effects of various form, up to the exact shape of the truncated-Gaussian and uniform distributions used to generate the exposed distributions. The explicit form of each model is reported in table 1. All models were fitted by minimizing the negative log-likelihood of the data given the parameters, using a quasi-Newton optimization algorithm (MATLAB’s *fminunc* function). For each model, the Bayesian information criterion (BIC; Schwarz, 1978) was computed from the log-likelihood. Model comparison was performed for the two types of exposed distributions separately, by subtracting all BIC values from the largest one. Differences in BIC (ΔBIC) larger than 2 are considered positive evidence against the model with the higher BIC.

**Table 1.**
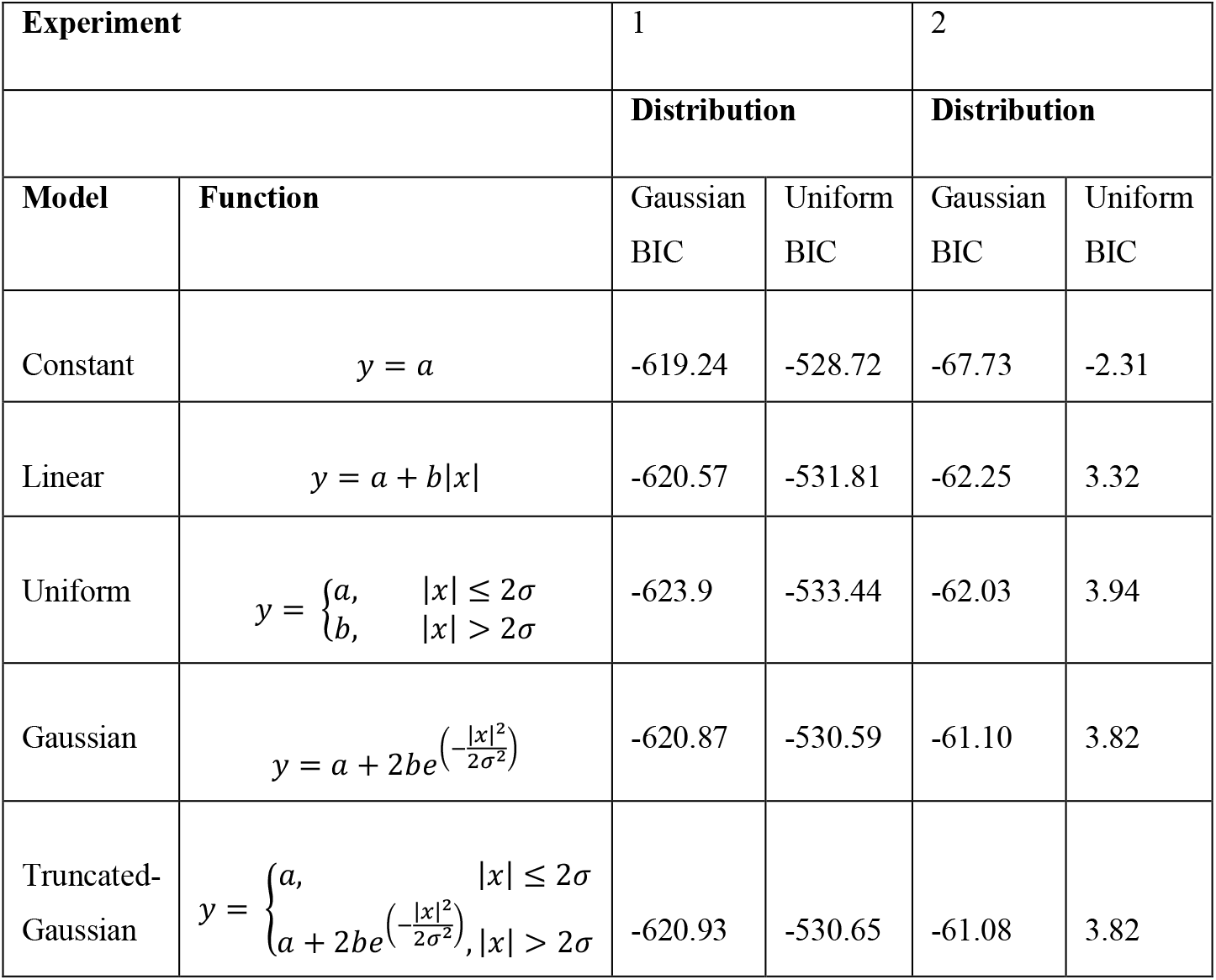
The set of models used in both experiments and the BIC of each model for trials with uniform and Gaussian distributions presented during exposure.

Model comparison was paralleled with a two-way repeated-measures ANOVA with factors: Distribution Type (2 levels) and CT-PD (6 levels, from 0° to 75° in steps of 15°, considering the absolute value of the original CT-PD levels). For the ANOVA analysis, the response times of each participant were averaged across CT-PD levels. In the location priming analysis of Experiment 1, we compared response times as a function of whether the location of the singleton in the last exposed display was the same or different from the location of the test singleton, using paired t-test analysis.

## Results

### Experiment 1

In experiment 1, where observers passively viewed arrays of orientated lines where one item was an orientation singleton, there was strong learning of the orientation distributions beyond the average or range. But there was little evidence that observers learned differences between the Gaussian and uniform distributions.

To evaluate whether the orientation distribution in the exposure displays affected performance on the test trial, we compared how well a set of models fit the shape of response times (RT) as a function of CT-PD (see Table 1). The set of models included a model with only a constant (i.e., no effect of the prior distractor distribution), a linear and a Gaussian model, and a uniform and a truncated-Gaussian model that resembled those used in generating the original distributions of the exposed orientations (see Figure 2A and Table 1).

**Figure 2.**
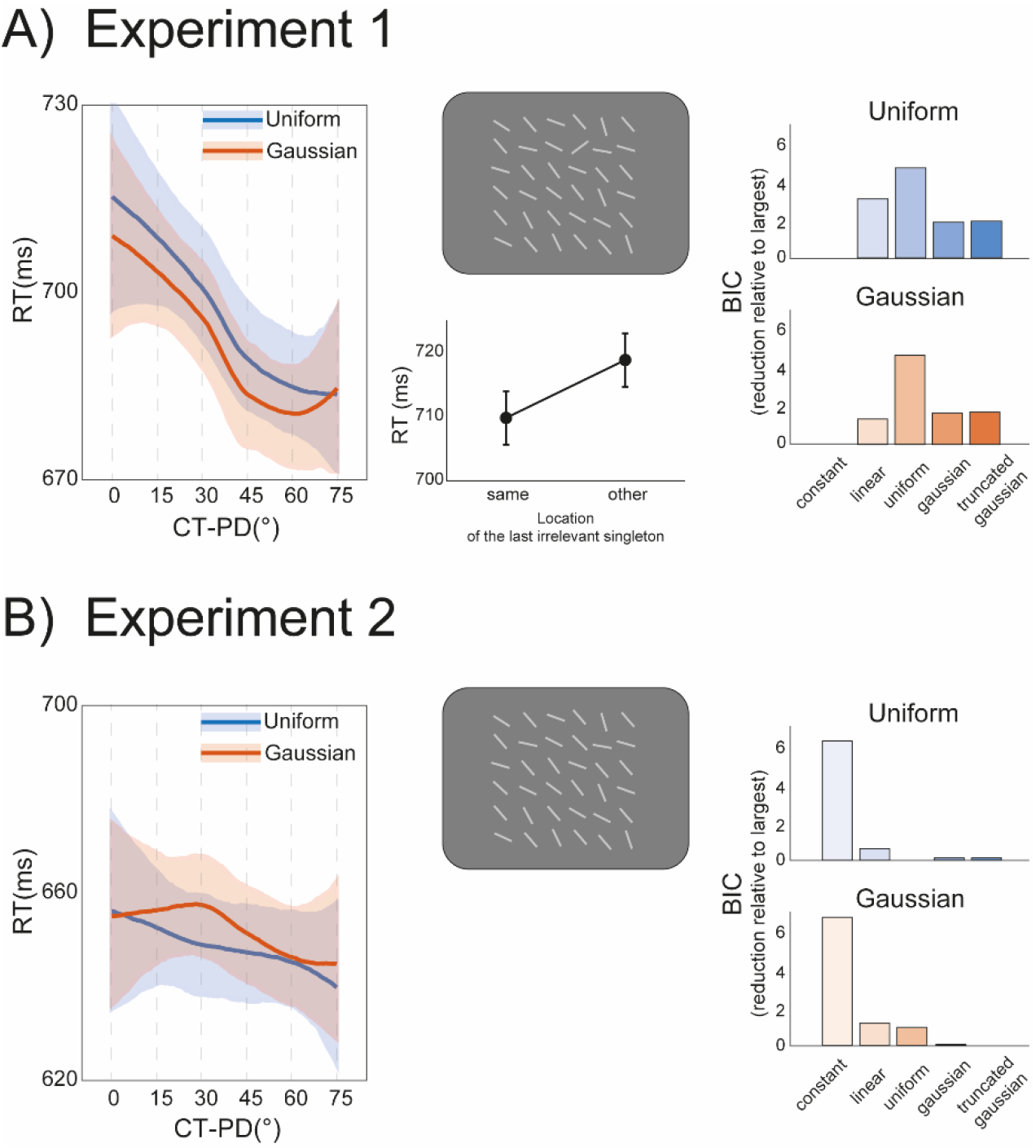
Results of Experiment 1 and 2. A) In Experiment 1, the exposed set of distractors contained a singleton orientation (as in the example display). Despite the absence of an explicit task, participants learned the distractors distribution during exposure, as evident from the decrease of response times (RT) as the angular distance between the previous distractors mean and the current target orientation increased (current target – previous distractor, CT-PD), a typical hallmark of FDL. According to Bayesian Information Criterion analysis (BIC), all models assuming an effect of the exposed distractors distribution performed better than a model assuming no effect (right panel, see also Table 1). This pattern was comparable across exposure sequences in which a uniform (blue bars) and a Gaussian distribution (red bars) were presented. The singleton line in Experiment 1 captured attention, causing significant negative location priming effects (response times were slower when the test target appeared at the same location of the last exposed singleton, line plot in the bottom-central panel). This involuntarily attentional capture likely caused the implicit learning of the distractors’ distribution. B) In Experiment 2, in which the singleton line was removed during exposure (as in the example display), FDL did not occur and a model assuming no effect of the previous distractor distribution performed better than all the other models (BIC plot, right panel).

All models assuming an effect of the prior distractor distribution performed better than the model assuming no effect (see Table 1; minimum ΔBIC across model comparisons against the constant = 1.32). In particular, model comparison through the Bayesian Information Criterion (BIC; Schwarz, 1978), revealed positive evidence favoring the uniform over the constant and over the other models (all ΔBIC > 2, except for the comparison ‘uniform vs. linear’ in the condition with uniform distractor distributions: ΔBIC = 1.62).

This pattern was similar for the two types of distributions observers were exposed to during the learning trials. This means that observers’ search times were clearly affected by the distribution of orientations during the exposure stage, a result further validated by a significant main effect of CT-PD, in a two-way repeated-measures ANOVA with the factors Distribution Type X CT-PD (main effect of CT-PD: F(5,115) = 6.86, *p* < 0.001, 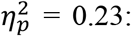 main effect of Distractor Type and interaction not significant). These results show that observers can passively learn the characteristics of a distribution of features, beyond the simple average. However, the learned characteristics did not allow sufficient resolution to distinguish between the shape of the uniform and the Gaussian distributions.

One potential explanation for this form of learning is that the presence of a singleton, although not task-relevant, per se, might have triggered automatic attentional capture mechanisms that lead to the implicit learning of the distractor features. That is, the singleton may have captured attention because it was an outlier in the orientation distribution and was implicitly represented and learned. To evaluate this possibility, we inspected potential priming effects due to the location of the irrelevant singleton in the last exposed display before the test trial. A small but significant negative priming effect revealed that search times were slower when the singleton during the last learning trials was in the same quadrant as the target singleton (t(23) = −2.24, *p* = 0.035). This confirmed that observers were not completely immune to the presence of a singleton at the exposure stage, and therefore, FDL might have been due to the automatic segmentation of a singleton from the distribution of features. In other words, the feature contrast causes singleton detection which is, in turn also connected with the learning of distributions. To address this point more directly, we performed experiment 2 in which no singleton appeared during the exposure stage.

### Experiment 2

As in the first experiment, observers passively viewed the display during the exposure trials and were tested on a single test trial. No evidence of an effect of the exposed distributions from the learning trials was found in this experiment (see Figure 2B and Table 1), with all the models performing worse than the constant only model (constant model against each of the other models with higher BIC, all ΔBIC > 4), and no significant main effect or interaction in the two-way repeated-measures ANOVA with factors Distribution Type X CT-PD (all *p* > 0.05). This means that the presence of a singleton during the learning trials in Experiment 1, was crucial for triggering FDL by passive viewing.

## Discussion

Recent work has shown that human observers can rapidly learn rich and detailed representations of distributions of visual features (Chetverikov et al., 2016, 2017a), above what is proposed in summary statistics accounts (Cohen et al., 2016). Here we asked whether this form of learning, in which the attentional system represents the full shape of a distribution of visual features, occurs during passive viewing or whether it requires active visual search.

We found that FDL occurs after passive exposure and in the absence of an explicit search task, provided that the passively viewed displays contain a singleton element. In previous work, observers also performed a singleton search in the learning trials, and the RT patterns following learning tracked the precise shape of the learned distributions. It was possible to distinguish, for instance, learned representations that resembled the shape of Gaussian, uniform, and even more complex distributions (Chetverikov et al., 2016, 2017b, 2019). In Experiment 1, observers’ performance was affected by the distribution of features during the exposure stage, beyond the simple average, but the resulting pattern of RT did not allow us to distinguish between the Gaussian and the uniform distributions.

One explanation for the inability to distinguish different distribution types is that this form of learning is eventually shaped and refined by the repeated attentional selection of a target singleton during active search. The detection of a deviant ‘outlier’ during singleton search may indeed be mediated by an implicit representation of the statistical properties of the whole display (Haberman & Whitney, 2012). Once the active process of segmenting out a deviant from the same distribution of features is reiterated a few times, the shape of the distribution is learned and temporarily stored in detail. Several findings reinforce this idea, suggesting that statistical representations are the building blocks of segmentation and categorization processes (Im et al., 2021; Khayat & Hochstein, 2018; Utochkin, 2015). In line with this, the most likely explanation for our results is that the irrelevant singleton on passive trials triggered an unsolicited and involuntary attentional capture, which in turn, led to the automatic segmentation of a deviant from the distribution, and a coarse representation of the distribution’s shape. This is supported by the absence of learning after removing the singleton from the passive trials of Experiment 2, and even more, by the negative location priming effect found in Experiment 1. The negative location priming suggests that participants did encode the irrelevant singleton (and its location) during passive viewing. This, in turn, impaired performance when the target appeared in the same location as the previous singleton, a form of negative priming typically observed at the location of previously irrelevant and distracting stimuli (Fox, 1995). We therefore speculate that involuntary detection of a deviant singleton left a coarse trace of the statistical distribution of visual features from which the singleton was an outlier. In more active conditions, where the singleton is the target of a visual search, the characteristics of the distribution may be learned and retained more in detail (Chetverikov et al., 2016).

Our results add to the emerging field of research showing latent forms of short-term plasticity in the attentional system. These studies have provided evidence of basic and non-associative forms of learning, like habituation of the attentional capture response, after passive and task-free exposure to single stimuli (Turatto et al., 2018; Turatto & Pascucci, 2016; Won & Geng, 2020). Here we show that involuntary attentional capture and singleton segmentation can foster more complex forms of learning in which the properties of a distribution of visual features are latently and automatically learned. The dependence of passive FDL on the presence of an outlier might be functionally meaningful: constantly learning the entire statistics of irrelevant features in the visual world might be redundant and resource-consuming; learning the distribution of features that signal a ‘surprise’, can aid the attentional system in directing resources more efficiently when such surprise becomes relevant for behavior.

## Data availability

All data will be made available in online repositories upon acceptance.

## Authors contribution

All authors contributed to the study idea and design. D.P. coded the experiment and main analysis. G.C. collected the data. D.P. drafted the initial manuscript. All authors contributed to the revision and final version of the manuscript.

## Aknowledgments

This research was supported by funding from the Swiss National Science Foundation (grant no. PZ00P1_179988 to DP), from the Icelandic Research Fund (#207045-052 to AK & DP) and the Research Fund of the University of Iceland (to AK). The funders had no role in study design, data collection and analysis.

